# The neurohormone tyramine stimulates the secretion of an Insulin-Like Peptide from the intestine to modulate the systemic stress response in C. elegans

**DOI:** 10.1101/2024.02.06.579207

**Authors:** Tania Veuthey, Sebastián Giunti, María José De Rosa, Mark Alkema, Diego Rayes

## Abstract

The DAF-2/insulin/insulin-like growth factor signaling (IIS) pathway plays an evolutionarily conserved role in regulating reproductive development, lifespan, and stress resistance. In *C. elegans*, DAF-2/IIS signaling is modulated by an extensive array of insulin-like peptides (ILPs) with diverse spatial and temporal expression patterns. However, the release dynamics and specific functions of these ILPs in adapting to different environmental conditions remain poorly understood. Here, we show that the ILP, INS-3, plays a crucial role in modulating the response to different types of stressors in *C. elegans*. *ins-3* mutants display increased resistance to both heat and oxidative stress; however, under favorable conditions, this advantage is countered by slower reproductive development. *ins-3* expression in both neurons and the intestine is downregulated in response to environmental stressors. Conversely, the neurohormone tyramine, which is released during the acute flight response, triggers an upregulation in *ins-3* expression. Moreover, we found that tyramine negatively impacts environmental stress resistance by stimulating the release of INS-3 from the intestine. The subsequent release of INS-3 systemically activates the DAF-2 pathway, resulting in the inhibition of cytoprotective mechanisms mediated by DAF-16/FOXO and HSF-1. These studies offer mechanistic insights into the brain-gut communication pathway that weighs adaptive strategies to respond to acute and long-term stress scenarios.

## INTRODUCTION

Organisms are constantly confronted with environmental challenges that require adaptive responses for their survival and reproductive success. These adaptive responses are orchestrated through sophisticated regulatory mechanisms that control a spectrum of physiological adjustments [1,2]. For example, when conditions are optimal, i.e. abundant food resources and favorable temperatures, organisms tend to prioritize and expedite their growth and reproductive development to provide them with a competitive advantage [3,4]. Conversely, in the face of adverse environmental conditions, such as a shortage of food or oxidative stress, animals adopt a contrasting strategy in which they slow down their growth and reproductive rate and redirect their energy resources towards cytoprotection [5,6]. The evolutionarily conserved insulin/insulin-like growth factor signaling (IIS) pathway plays a pivotal role in this strategic plasticity of animals [7]. It is well-established that elevated insulin levels are associated with cell growth and development [7,8], whereas reduced insulin levels are linked to increased lifespan and greater resistance to environmental long-lasting stressors such as high temperature or oxidation [9–11]. The mechanisms by which the release of Insulin-Like Peptides (ILPs) is coordinated in response to environmental conditions remain mostly unknown across all animals.

The complexity of mammalian stress physiology makes it challenging to investigate the fundamental mechanisms governing the systemic control of cellular defense processes. Genetic studies in the nematode *C. elegans* have played a crucial role in unveiling the significance of the conserved IIS pathway in controlling both lifespan and stress resistance [11–13].

Although *C. elegans* has a single insulin receptor, DAF-2, the worm’s genome encodes 40 ILPs that can interact with this receptor [14]. The downregulation of DAF-2 typically extends lifespan and enhances resistance to environmental stressors [12]. ILPs have been reported to modulate aversive olfactory learning [15], lifespan, dauer formation [14,16,17], and germ cell proliferation through the IIS pathway [18]. However, no mutation or deletion of single insulin genes has been found to fully replicate the phenotypes observed in the absence of DAF-2 [19]. Furthermore, the overexpression of each of the forty insulin genes revealed that these ILPs can act not only as strong agonists but also as weak agonists, antagonists, or even exhibit pleiotropic functions on the DAF-2/IIS pathway [20]. Therefore, deciphering the specific roles of individual ILPs is challenging. Just as in most animals, where ILPs act as hormones, in *C. elegans*, they are secreted into the pseudocoelom, the fluid-filled body cavity, acting as long-range signaling molecules [7]. Importantly, the mechanisms underlying ILPs secretion in *C. elegans* shares similarities with mammals, and conserved factors such as ASNA-1, HID-1, and GON-1 have been identified to play a role [21–23].

Under acute life-threatening situations, such as imminent predation or aggression, organisms trigger a high-energy demanding, short-lived “flight response” in a bid to enhance their odds of escape [24–26]. We have previously uncovered a novel brain-gut communication pathway in *C. elegans* where neural stress hormones released during the flight response negatively impact health by activating the DAF-2/IIS pathway [27]. The flight response triggers the activation of a specific pair of neurons that release tyramine, the invertebrate equivalent of adrenaline [27,28]. Tyramine coordinates different motor programs of the flight response, allowing the worm to escape from predation [29–32]. However, the downregulation of tyramine signaling is essential to deal with environmental stressors when a behavioral response alone fails to alleviate the exposure to a threatening stimulus [27]. Specifically, we found that tyramine release during the flight response activates an adrenergic-like receptor, TYRA-3, in the intestine. TYRA-3 activation subsequently leads to the stimulation of DAF-2/IIS pathway throughout the body. Systemic activation of the DAF-2/IIS pathway may increase the metabolic rate required for the physical and energy demands of the flight response but it comes at a cost. DAF-2/IIS pathway stimulation inhibits cytoprotective mechanisms that are needed to cope with longer lasting environmental stressors. Thus, tyramine acts as a state-dependent neural signal that controls the switch between acute and long-term stress responses. The processes linking the tyraminergic activation of TYRA-3 in the intestine with the systemic stimulation of the DAF-2 pathway remains unknown.

Here, we find that animals mutant for the Insulin-Like Peptide INS-3, display an increased resistance to environmental stressors such as oxidation and heat. We reveal that the activation of the tyraminergic receptor TYRA-3 induces the release of INS-3 from the intestine. In the context of a sustained escape response, the tyramine-induced release of INS-3 is crucial for the systemic activation of the DAF-2 pathway, leading to the suppression of cytoprotective mechanisms dependent on DAF-16/FOXO and HSF-1. This renders these animals more vulnerable to environmental stressors. By elucidating the tyramine-mediated secretion of INS-3, we unveil the complete molecular sequence underlying the detrimental effects of a sustained flight-stress response in *C. elegans*.

## RESULTS

### Intestinal expression of ins-3 decreases C. elegans resistance to environmental stressors

The *C. elegans* genome encodes an extensive family of ILP genes, with a substantial number of them expressed in the intestine [19]. Our previous work suggested that ILPs released from the intestine might play a key role in modulating the stress response [27]. To test this hypothesis we used RNA interference (RNAi) to silence the expression of intestinal ILPs that have been reported to act as strong agonist of DAF-2/IIS signaling pathway [20]. We examined resistance to the oxidizing agent FeSO_4_ (15 mM) in animals treated with *ins-3, ins-4, ins-6, ins-32 and daf-28*, RNAi. RNAi silencing of *ins-3* increased oxidative stress resistance (Fig 1A). Similar to *ins-3* RNAi-silenced animals, null mutants in the *ins-3* gene are significantly more resistant to oxidative stress than control animals (Fig 1B). Furthermore, *ins-3* mutants are also more resistant to thermal stress (35°C, 4 hrs) than control animals (Fig 1B).

**Fig 1.**
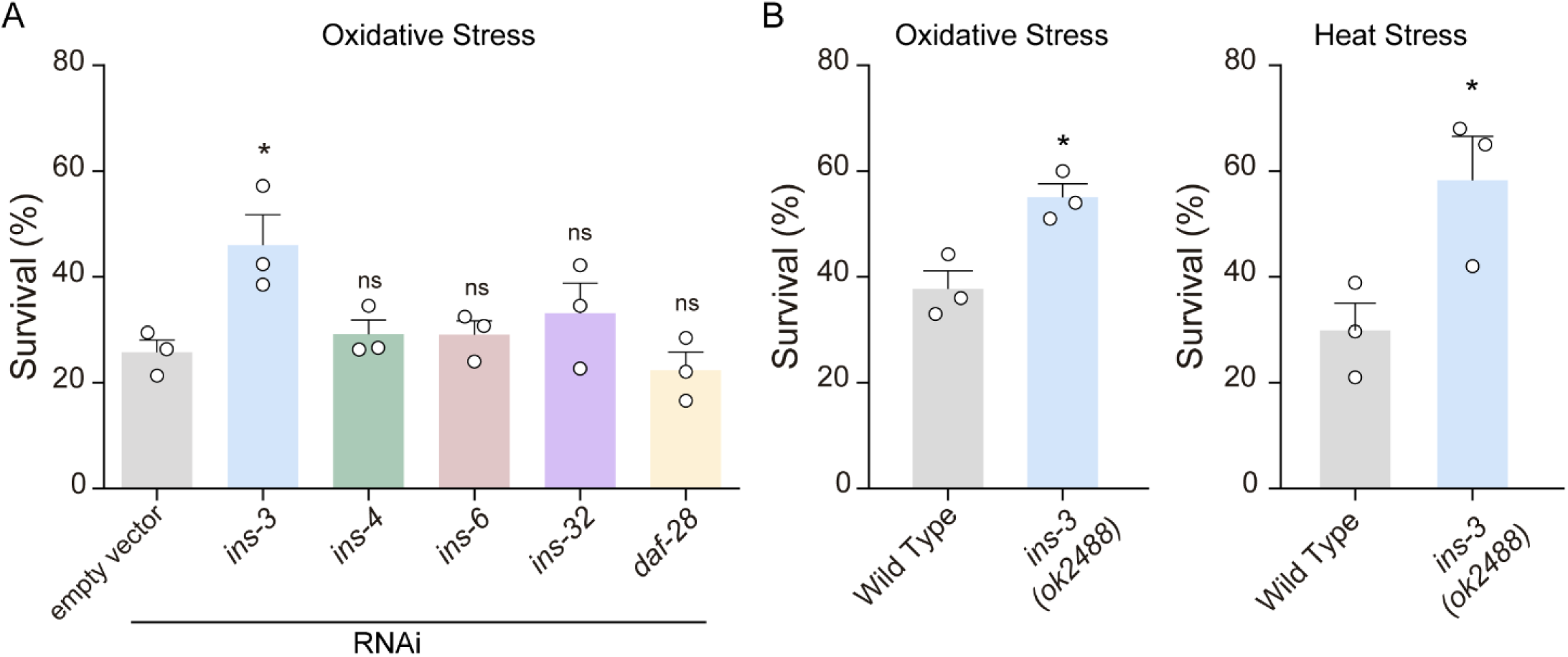
The deficiency of the Insulin-Like Peptide INS-3 confers enhanced resistance to environmental stressors. A) Survival percentages in response to oxidative stress (1 h, 15 mM FeSO4) in animals subjected to RNAi-mediated silencing of Insulin-Like Peptides that are expressed in the intestine, and were previously identified as potent agonists of the DAF-2 receptor. Mean ± s.e.m., n = 3, 60–80 animals per condition per experiment. One-way ANOVA, Holm–Sidak’s post-hoc test for multiple comparisons. ns: not significant, * p<0.05. B) Survival percentages of wild type and *ins-3* null mutant worms exposed to oxidation (1h, 15 mM Fe SO4) and heat stress (4 hrs at 35°C). Two-tailed t-test was used, n = 3, 40–80 animals per condition per experiment. * p<0.05

Using a *Pins-3::GFP* transcriptional reporter we found that *ins-3* is mainly expressed in the intestine and a subset of neurons consistent with previous reports (Fig 2A) [19]. Transgenic expression of *ins-3* under control of its endogenous promoter in a *ins-3* null mutant background restored oxidative stress sensitivity to wild type levels (Fig 2B). To determine where *ins-3* acts to modulate the stress response, we performed tissue-specific rescue experiments. Expression of *ins-3* in the intestine but not in neurons, completely restored oxidative and heat stress sensitivity of *ins-3* null mutants to wild type levels (Fig 2B). These findings indicate that INS-3 release from the intestine inhibits the animaĺs capacity to cope with environmental stressors.

**Fig 2.**
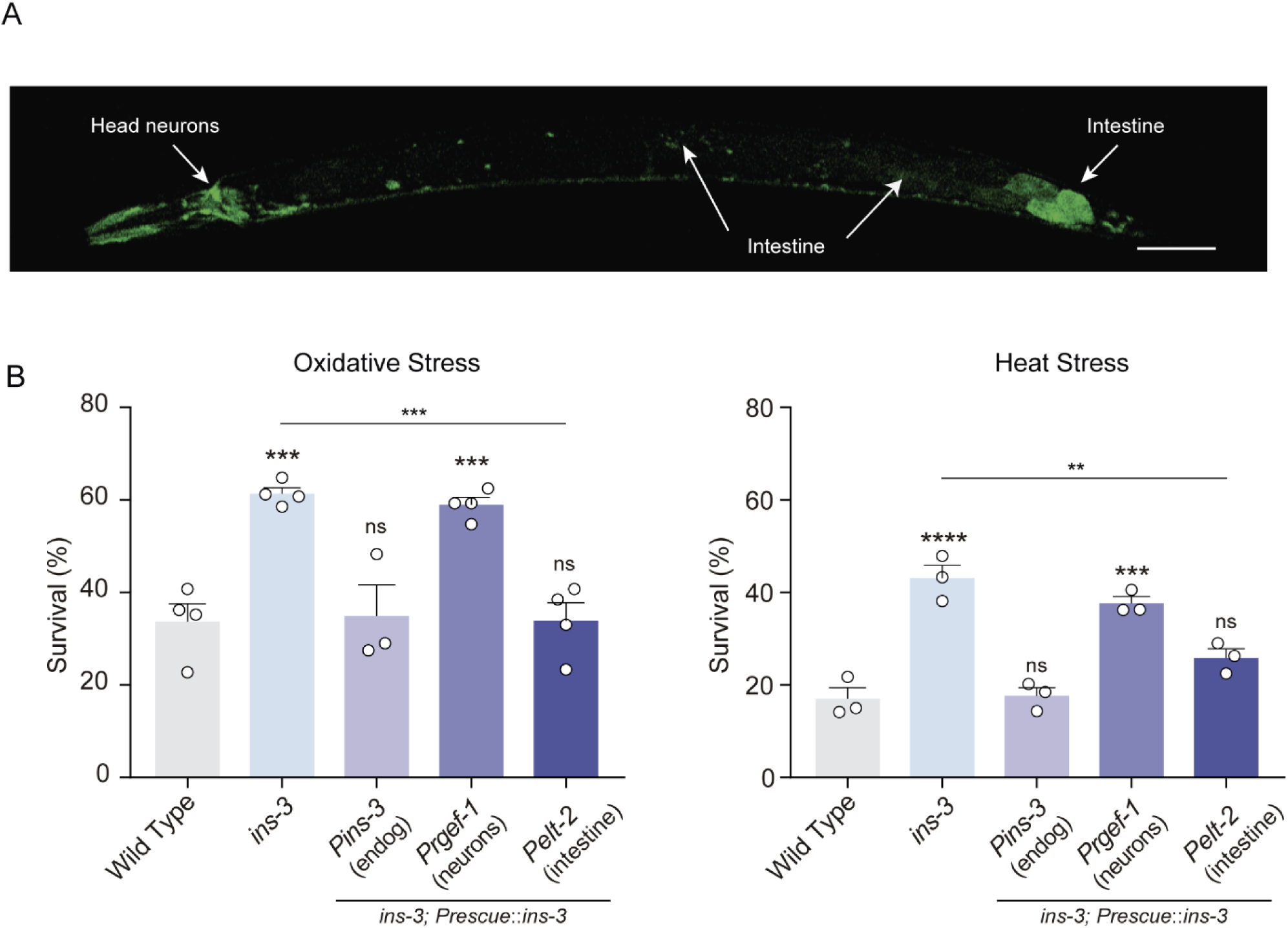
INS-3 in the intestine modulates the stress response. A) Expression of *Pins-3::GFP* reporter. Scale bar 50 µm B) Stress resistance of *ins-3* mutant animals expressing an *ins-3* cDNA driven by *Pins-3* (endogenous), *Prgef-1* (pan-neuronal) or *Pelt-2* (intestinal) promoters upon exposure to oxidative stress (left) or heat stress (right). Mean ± s.e.m., n = 3-4, 40-60 animals per condition per experiment. One-way ANOVA, Holm–Sidak’s post-hoc test for multiple comparisons was used. Two-tailed t-test for comparison with *ins-3* null mutant was used. ns: not significant, ** p<0.01 *** p<0.001, **** p<0.0001.

### INS-3 is a systemic agonist of DAF-2/IIS signaling pathway

Activation of DAF-2 typically results in the phosphorylation of DAF-16 and its retention in the cytosol [9]. To investigate whether intestinal INS-3 is systemically activating the DAF-2 pathway, we assessed the subcellular localization of the transcription factor DAF-16, using the translational *Pdaf-16::daf-16::GFP* reporter. Under basal conditions, DAF-16 primarily localized to the cytoplasm in both wild type and *ins-3* null mutant backgrounds (Fig S1A). We have previously shown that, upon exposure to heat (35°C), the nuclear localization of DAF-16 in wild type animals increases, reaching its maximum after 30 minutes [27]. After mild stress induction (10 min at 35°C), *ins-3* null mutants showed markedly enhanced nuclear accumulation of DAF-16 compared to the wild type (Fig 3A). Importantly, tissue specific rescue of *ins-3* in the intestine decreased nuclear DAF-16 accumulation (Fig 3A).

**Fig 3.**
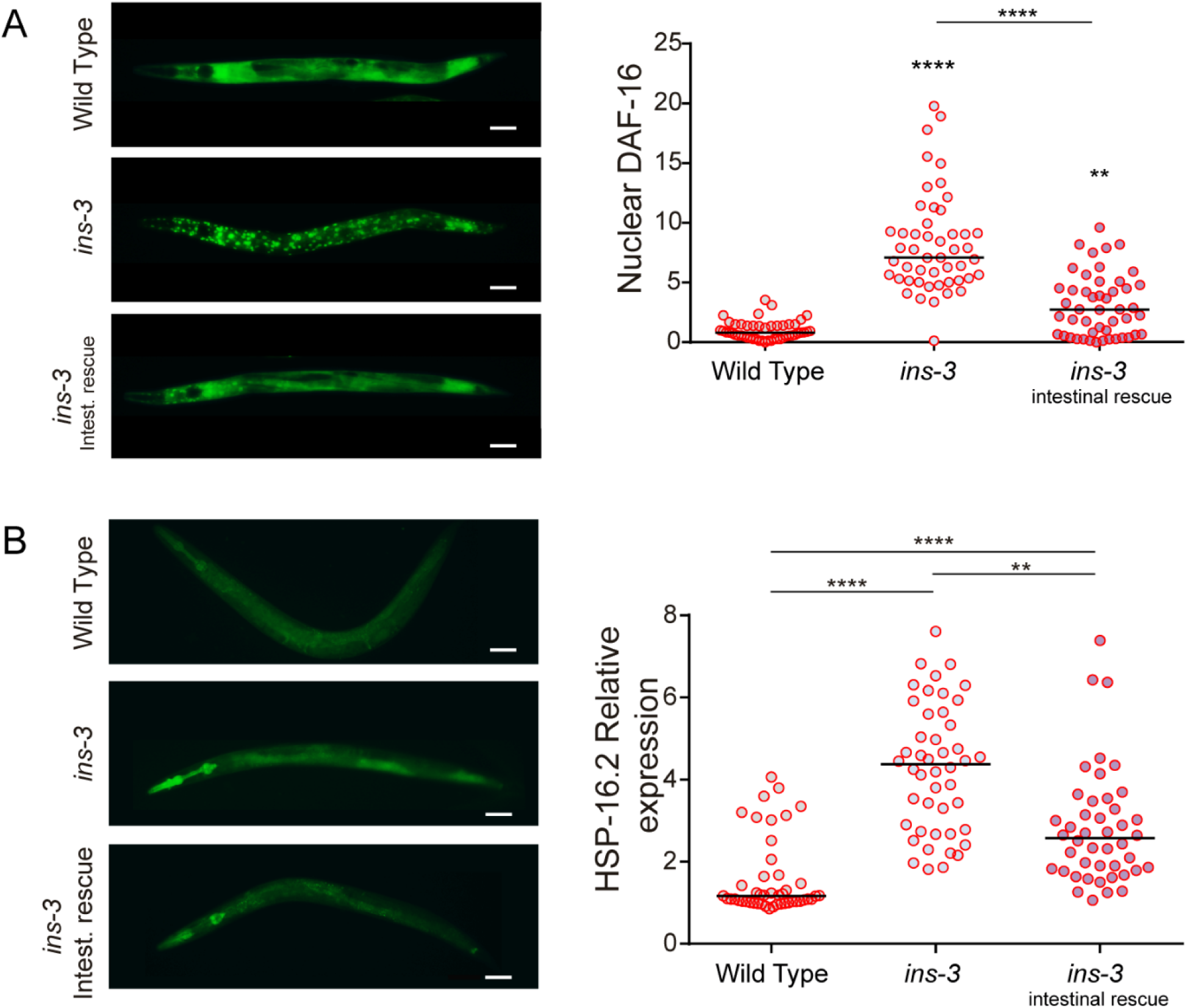
INS-3 constrains DAF-16/FOXO and HSF-1 dependent cytoprotective defensive mechanisms. A) Left: Representative fluorescence images (20 X) depicting the localization of DAF-16a/b::GFP after a short heat exposure (35°C for 10 minutes) in WT, *ins-3* null mutant, and intestinal rescue of *ins-3* backgrounds. Scale bar, 50 μm. Right: Corresponding scatter dot plots with the number of cells with nuclear DAF-16 per animal (normalized to naive animals, line shows median). n= 40-50 per condition. One-way ANOVA with Dunn’s multiple comparisons post-hoc test. ** p<0.01**** p<0.0001 B) Left. Representative fluorescence images (20 X) of animals expressing *Phsp16.2::GFP* in WT, *ins-3* null mutant, and intestinal rescue of *ins-3* backgrounds after 15 min of heat exposure (35°C) followed by 70 min recovery at 20°C. Scale bar, 50 μm. Right. Corresponding quantification of the fluorescence level per animal. Scatter dot plot with relative expression of *Phsp16.2::GFP* normalized to naive animals (line shows median). n= 40-50 per condition. One-way ANOVA and Dunnett’s post-hoc test versus naïve. ** p<0.01 **** p<0.0001

Activation of DAF-2 not only inhibits DAF-16 translocation but also leads to the suppression of other cytoprotective transcription factors, such as HSF-1 (Heat Shock Factor-1) [33]. To assess the activity of HSF-1, we analyzed the expression levels of *HSP-16.2*, an HSF-1 effector gene, using the transcriptional fluorescent reporter *hps-16.2::GFP*. We have previously demonstrated that *hsp-16* expression is induced by heat, much like the nuclear translocation of DAF-16 [27]. We found that, similar to our observation on DAF-16 translocation, under basal conditions *hsp-16.2*::*GFP* fluorescence intensity was similar between wild type and *ins-3* mutant animals (Fig S1B). However, after mild heat stress (15 min at 35°C), *ins-3* mutants exhibit increased *hsp-16.2*::*GFP* expression compared to wild type animals (Fig 3B). Intestinal *ins-3* expression decreased *hsp-16.2* expression of *ins-3* mutants, albeit not to wild type levels (Fig 3B). These results indicate that *ins-3* mutant animals trigger cytoprotective mechanisms more readily when exposed to environmental stressors. In addition, we found that *ins-3* mutants, produced a smaller number of offspring and exhibited slightly slower development compared to wild type animals (Fig S2), consistent with the down regulation of the DAF-2/IIS pathway [34]. Together our data show that INS-3 functions as an agonist in the regulation of the DAF-2/IIS pathway.

### ins-3 expression is differently modulated by distinct types of stressors

Exposure to different stressors can exert both positive and negative effects on the expression of ILPs [19,35]. To investigate whether stressors impact *ins-3* expression, we exposed animals expressing *Pins-3*::*GFP* to oxidative and thermal stressors (Fig 4). Exposure to either the oxidizing agent FeSO_4_ (1h) or heat (6 hrs at 30°C), led to a decrease in GFP expression in both the intestine and neurons (Fig 4). This decrease in fluorescence upon exposure to these stressors does not appear to be a non-specific effect on protein expression and/or stability, as the expression of another ILP, *ins-4*, was unaffected and even slightly increased upon oxidative stress (Fig S3).

**Fig 4.**
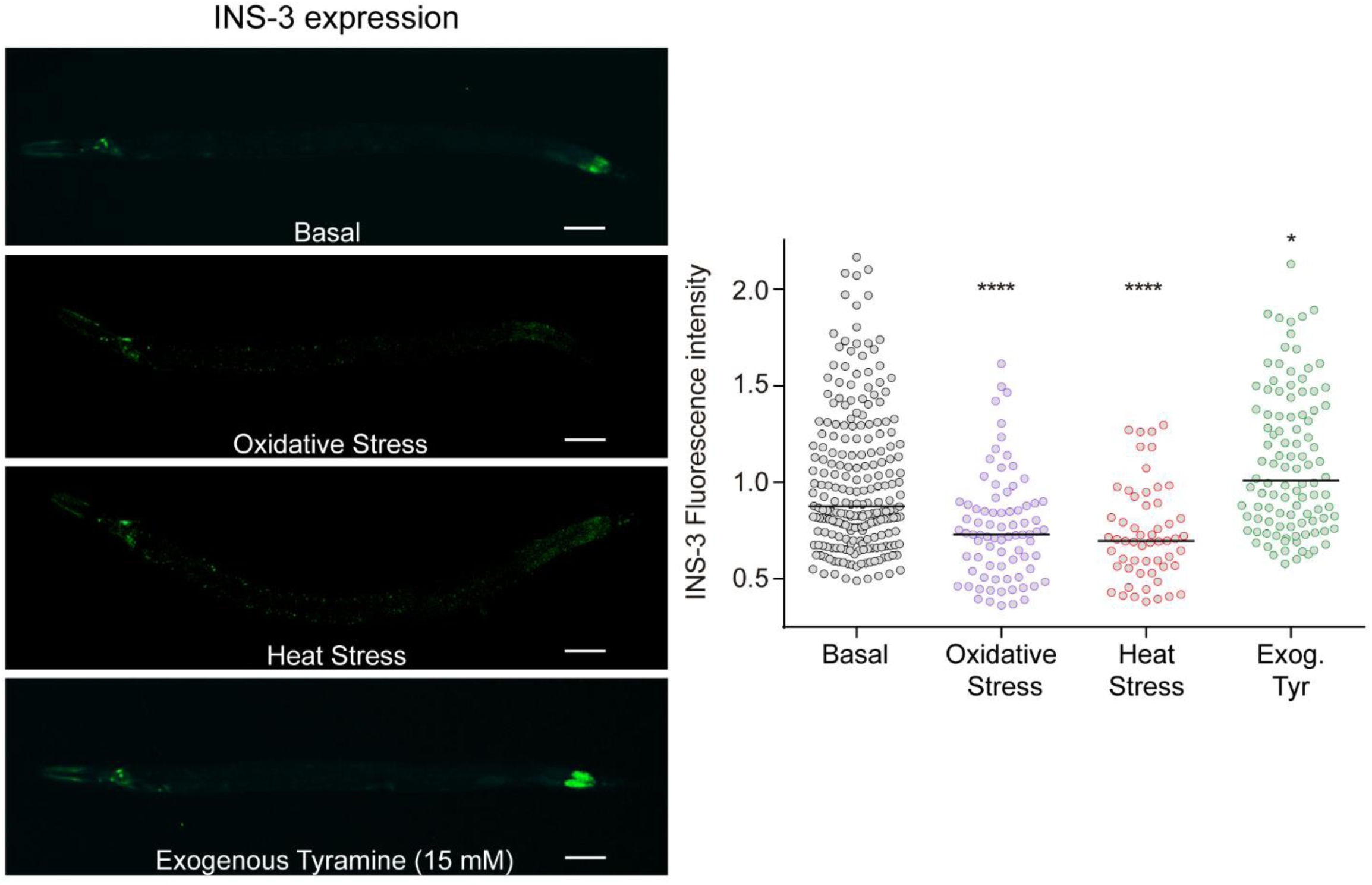
Environmental stressors and tyramine differentially modulate *ins-3* expression. Left: Representative epifluorescence images (20 X) of L4 animals expressing *Pins-3::GFP* under basal conditions (20°C on NGM plates seeded with OP50), under oxidative stress (1 mM FeSO4 for 2 hrs) or heat stress (30°C for 6 hrs) and in the presence of exogenous Tyramine (15 mM). Scale bar 50 µm. Right. Corresponding quantification of fluorescence levels per worm. Scatter dot plot with relative expression of *Pins-3::GFP* normalized to naive animals. Line at the median. n= 50-200 per condition. One-way ANOVA and Dunn’s post-hoc test versus basal was used. * p<0.05 *** p<0.001

We have previously shown that exposure to environmental stressors such as oxidation or heat leads to a downregulation of tyramine signaling [27]. Animals unable to downregulate this signal become much more sensitive to environmental stressors due to systemic hyperactivation of the DAF-2/IIS pathway [27]. In contrast to environmental stressors, exposure to exogenous tyramine significantly enhances the expression of the transcriptional *ins-3*::*GFP* reporter in the intestine (Fig 4). Our data indicate that environmental stressors and the acute flight response have opposing effects on *ins-3* expression.

### Tyramine triggers INS-3 release from the intestine

The release of tyramine during the flight response stimulates the DAF-2/IIS signaling pathway response through the activation of the Gq-coupled TYRA-3 receptor in the intestine [27]. We hypothesized that TYRA-3 activation may trigger the release of agonist ILPs from the intestine. To determine whether tyramine stimulates the intestinal release of INS-3 from the intestine, we generated transgenic animals expressing the translational reporter *ins-3::venus* driven by the intestinal promoter *Pges-1*. Secreted Venus-tagged peptides are taken up by the coelomocytes, scavenger cells which continually endocytose material secreted into the pseudocoelomic fluid. Fluorescence intensity in the coelomocytes can be therefore used as a proxy for peptide secretion [21,36,37]. In naïve animals, we observed faint intestinal expression of *ins-3::venus*. Additionally, we detected fluorescence in coelomocytes, suggesting a tonic release of INS-3 under basal conditions (Fig 5; Fig S4). When *ins-3::venus* animals were exposed to 15 mM tyramine we observed a 2-fold increase in the fluorescence intensity of coelomocytes (Fig 5). The enhanced fluorescence is not a consequence of tyramine inducing an increase in *Pges*-1::*ins-3::venus* transcription, as we did not detect increase in intestinal fluorescence in animals treated with tyramine. Therefore, we conclude that tyramine stimulate the secretion of intestinal INS-3 (Fig 5).

**Fig 5.**
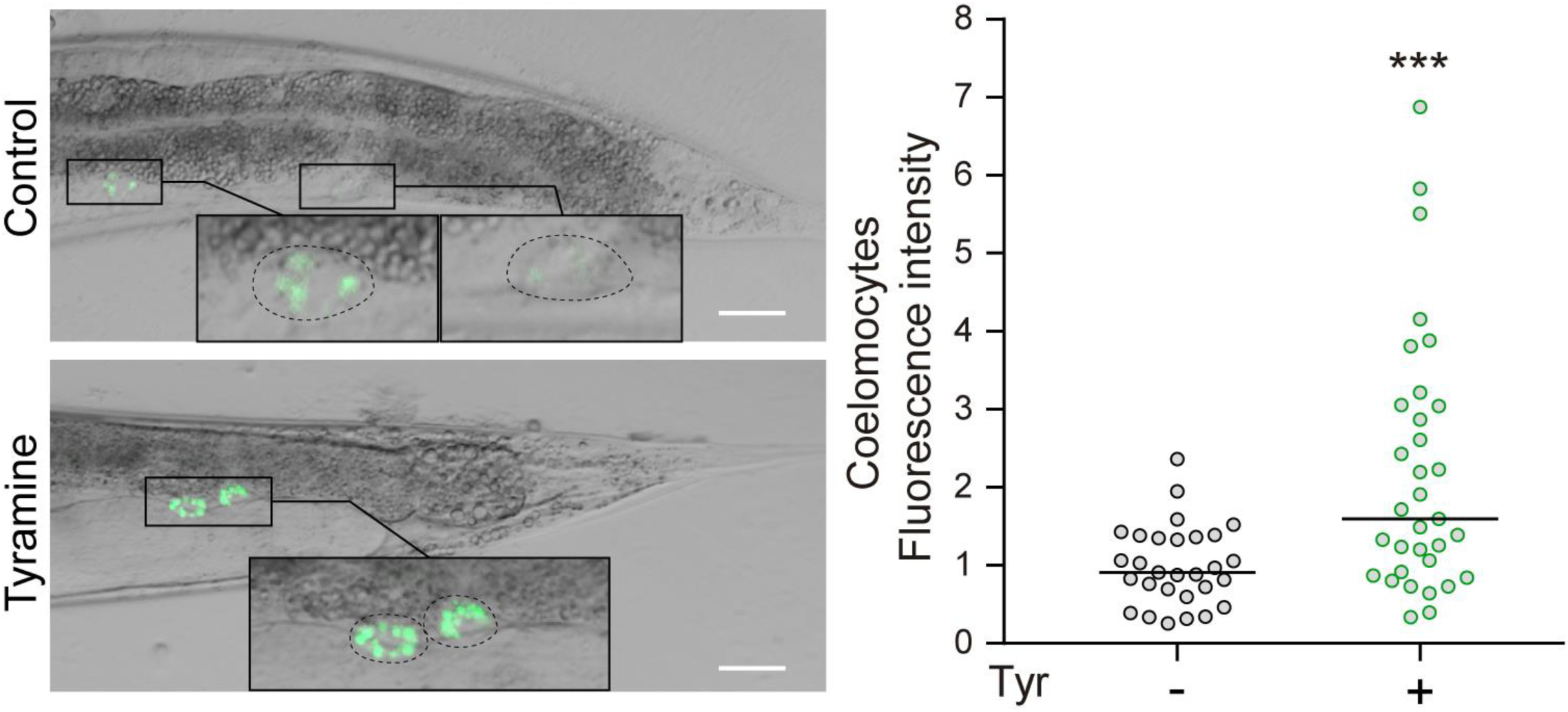
Tyramine triggers INS-3 release from the intestine. Left. Representative epifluorescence images (40 X) of young adults expressing *Pges-1::ins-3::venus* in the absence or presence of exogenous Tyramine (15 mM). Scale bar 25 µm. Fluorescence is located in the two posterior coelomocytes of worms (inset). Right. Corresponding quantification of fluorescence intensity on the coelomocytes (normalized to naive animals). Line at the median. n= 25-30 per condition. Two-side t-test was used. *** p<0.001

### Tyramine inhibits cytoprotective mechanisms through intestinal INS-3 secretion

Is INS-3 secretion required for the tyraminergic activation of the DAF-2/IIS pathway? To address this question, we examined the oxidative and thermal stress resistance of *ins-3* null mutants in the presence of exogenous tyramine. Unlike wild type animals, the addition of exogenous tyramine did not affect the stress resistance of *ins-3* null mutants (Fig 6A). Furthermore, when we selectively rescued *ins-3* expression in the intestine, the negative impact of exogenous tyramine on stress resistance is restored to wild type levels (Fig 6A).

**Fig 6.**
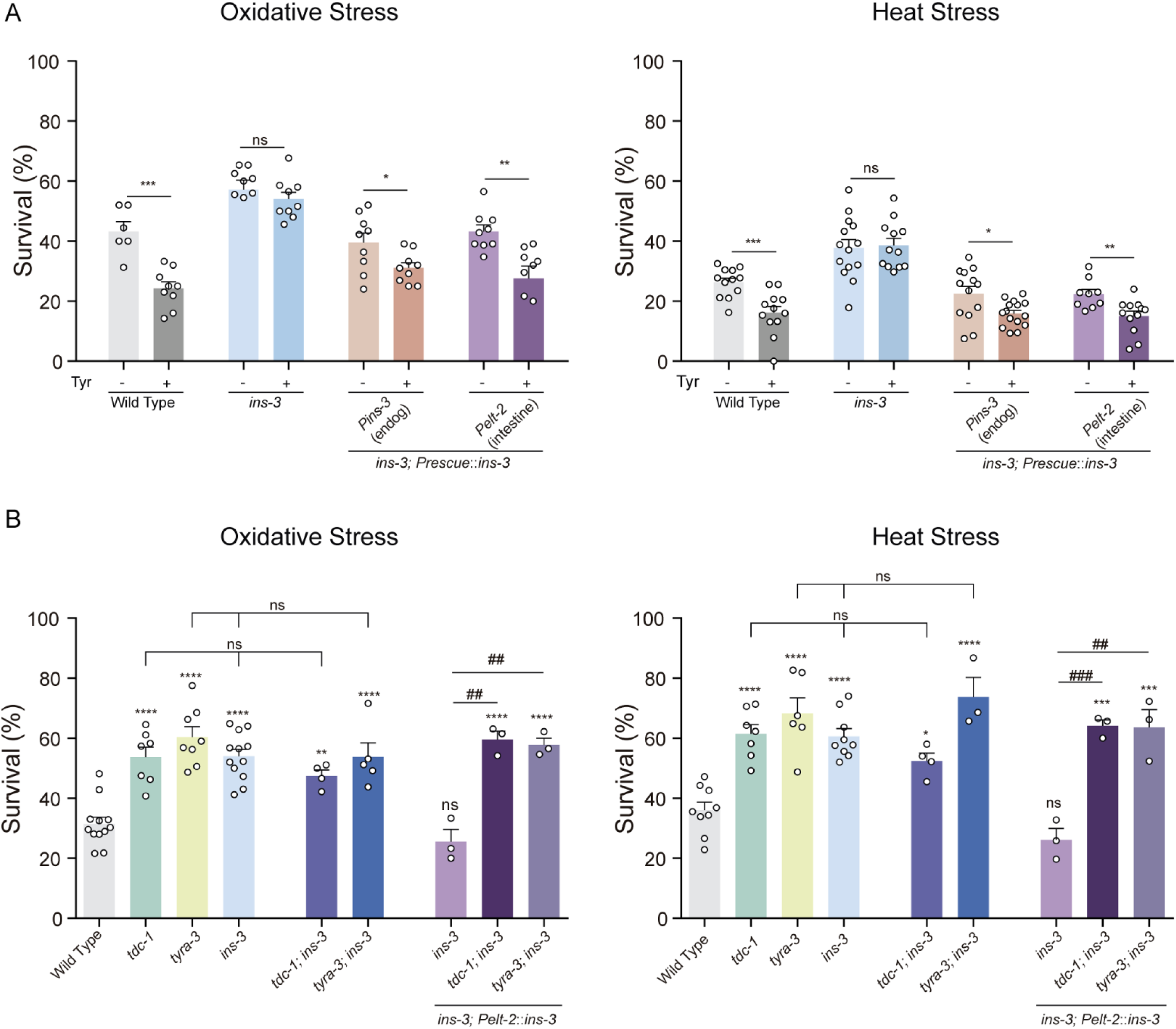
Tyramine and INS-3 act in the same pathway to modulate stress resistance. A) Stress resistance of *ins-3* mutant animals expressing an *ins-3* cDNA driven by *Pins-3* (endogenous) or *Pelt-2* (intestinal) promoters upon exposure to oxidative stress (left) or heat stress (right) in the absence or presence of exogenous Tyramine (15 mM). The detrimental effect of exogenous tyramine on stress resistance is only abolished on *ins-3* mutant animals. Mean ± s.e.m., n= 6-15 independent experiments. Each experiment included 40-60 animals per condition. For conditions with Tyr, two-tailed Student’s t-test versus same strain without Tyr was used. ns: not significant, * p<0,05 ** p<0.01, *** p<0.001. B) Survival percentages of animals exposed to oxidative (left) and heat stress (right). Results are shown as mean ± s.e.m., n= 3-12 independent experiments. Each experiments included 40-50 animals per condition. The stress resistance is not further enhanced in double mutants *tdc-1;ins-3* and *tyra-3;ins-3* compared to the single mutants. Additionally, the intestinal expression of *ins-3* does not restore resistance to wild-type levels in tyramine-deficient backgrounds. One-way ANOVA, Holm–Sidak’s post-hoc test versus wild type was used (*** p<0.001, **** p<0.0001). One-way ANOVA, Dunnett’s post-hoc test versus intestinal rescue of *ins-3* was used (## p<0.01, ### p<0.001). One-way ANOVA, Dunnett’s post-hoc test was used for mutants deficient in both tyramine and *ins-3* (*tdc-1;ins-3* and *tyra-3;ins-3*) compared to the corresponding single mutants. ns: not significant

We have previously shown that tyramine deficient, *tdc-1*, and the tyramine receptor, *tyra-3* mutants are resistant to both oxidative and thermal stressors [27]. We here found that the resistance of *ins-3* mutants is similar to that of *tdc-1* and *tyra-3,* mutants (Fig 6B). To further investigate the crosstalk between tyramine and INS-3 in modulating the stress response, we examined the resistance of *ins-3*; *tdc-1* and *ins-3; tyra-3* double mutants. These double mutants did not show any further enhancement in stress resistance compared to *tdc-1, ins-3* or *tyra-3* single mutants (Fig 6B). This observation is consistent with the notion that *tdc-1*, *tyra-3* and *ins-3* genes function in the same pathway to regulate the environmental stress response. Unlike our observations in *ins-3* null mutant backgrounds, the intestinal rescue of *ins-3* in *tyra-3;ins-3 and tdc-1;ins-3* backgrounds, did not impact the stress resistance (Fig 6B). Furthermore, exogenous tyramine failed to decrease stress resistance in the intestinal *ins-3* rescue strain in *tyra-3* mutant background (Fig S5). These findings strongly support the idea that INS-3 acts downstream of tyramine to modulate the stress response.

Since nuclear translocation of DAF-16 is enhanced in *ins-3* null mutant animals under mild stress (Fig 3A), we analyzed the effects of exogenous tyramine on DAF-16 localization in these mutants. After strong heat (30 min at 35°C), DAF-16/FOXO predominantly localized to the nucleus in both wild type and *ins-3* mutants (Fig 7A). Exogenous tyramine reduced DAF-16/FOXO nuclear localization in wild type animals, consistent with previous reports [27]. The addition of exogenous tyramine, however, failed to inhibit DAF-16 nuclear translocation in *ins-3* mutant animals (Fig 7A). Importantly, the expression of *ins-3* in the intestine reinstates the inhibitory effect of tyramine on the nuclear translocation of DAF-16/FOXO. This demonstrates that the negative modulation of DAF-16 by tyramine relies on the release of INS-3 from the intestine (Fig 7A). Taken together, our findings suggest that during the escape response the neurohormone tyramine leads to the release of INS-3 from the intestine, which, in turn, activates the DAF-2 pathway in various tissues, thus inhibiting cytoprotective mechanisms (Fig 7B).

**Fig 7.**
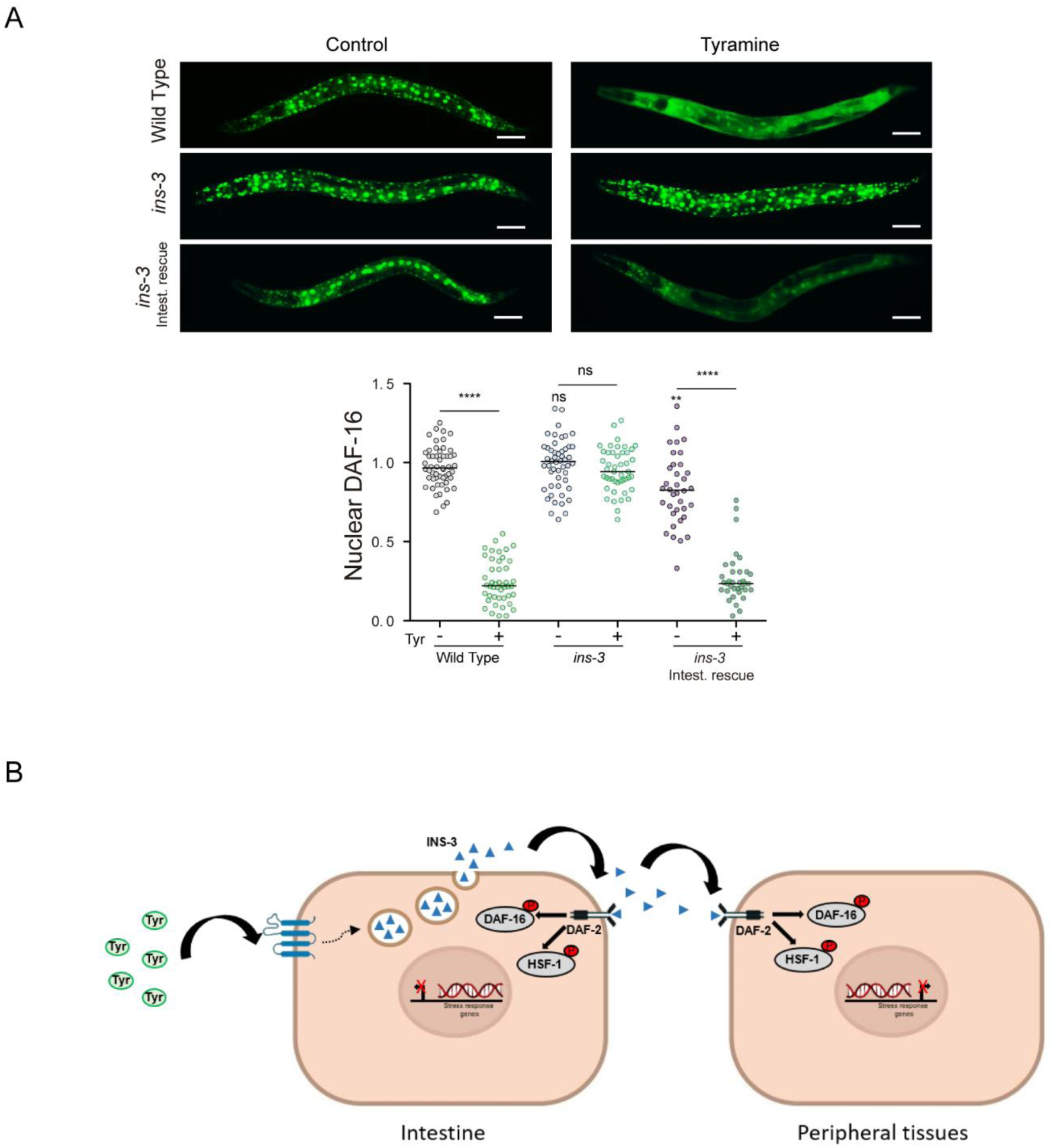
Tyramine inhibits cytoprotective mechanisms through intestinal INS-3 secretion. A) Top: DAF-16a/b::GFP localization upon strong heat (35°C) for 30 min, in the absence or presence of exogenous tyramine (15 mM Tyr, 30 min). Scale bar, 50 μm. Tyramine can inhibit DAF-16 nuclearization upon expression of *ins-3* in the intestine. Bottom: Corresponding scatter dot plots with the number of cells with nuclear DAF-16 per animal (normalized to naive animals, line shows median). n= 35-50 per condition. One-way ANOVA with Dunn’s multiple comparisons post-hoc test versus wild-type was used. ns: not significant, ** p<0.01. Two-tailed t-test (no Tyr versus Tyr) was used. ns: not significant, **** p<0.0001 B) Model: The escape neurohormone tyramine triggers the intestinal release of INS-3 to systemically activate DAF-2. DAF-2 activation leads to DAF-16 and HSF-1 phosphorylation precluding their nuclear translocation, therefore inhibiting the transcription of cytoprotective genes.

## DISCUSION

The insulin/IGF-1 signaling (IIS) pathway plays a crucial role in the stress response throughout the animal kingdom [7,38]. Previous studies have shown that downregulation of insulin signaling in *C. elegans* increases lifespan, and improves resistance to different environmental stressors [11,13,39–41]. While key components of the IIS pathway, such as the DAF-2 receptor and downstream molecules are well characterized, the specific roles of individual insulin-like peptides (ILPs) in the stress response remain largely unclear. The *C. elegans* genome contains 40 putative ILPs [42], with diverse temporal and spatial expression patterns with agonist or antagonist impact on the DAF-2 receptor, indicating a complex regulation of the DAF-2/IIS signaling pathway. Interestingly, most of these ILPs are expressed in the intestine [19]. Similar to other animals, the *C. elegans* intestine not only serves essential functions in digestion, but also plays pivotal role in the stress response [43]. Previous studies have shown that intestinal DAF-2/IIS is crucial for regulating resistance to oxidative stress, as intestinal-specific deletion of DAF-2 leads to increased oxidative stress resistance in a DAF-16-dependent manner [44]. In addition, the intestine also plays a non-cell autonomous role in the DAF-2-dependent control of lifespan [45–47].

We found that animals lacking the ILP INS-3 have an increased resistance to oxidative and heat stress. This occurs because INS-3 activates the DAF-2/IIS pathway, consequently inhibiting the response to environmental stress. While *ins-3* is expressed in the nervous system and the intestine of *C. elegans, ins-3* expression in the intestine was sufficient to restore oxidative and heat stress sensitivity to wild type levels, indicating that INS-3 release form the intestine modulates the stress response. Our findings strongly suggest that the intestinal ILP INS-3 plays a key role in modulating systemic environmental stress response, contributing to the broader network of stress regulation in *C. elegans.* Importantly, the production of ILPs in the intestine may not be unique to *C. elegans*, as the ability to express and secrete insulin by epithelial cells of the intestine has been reported in rats [48,49]. Furthermore, similar to our observations, this expression appears to be modulated by stressful conditions [50].

While there is increasing recognition of the importance of brain-gut communication in animal physiology and behavior, the molecular mechanisms underlying this interaction remain poorly understood. In previous work, we identified a novel brain-gut communication pathway that sheds light on how the repeated activation of the flight response can impair the reaction to other environmental challenges [27]. Specifically, we demonstrated that the release of tyramine from RIM neurons during the flight response activates TYRA-3 in the intestine. TYRA-3 activation, in turn, leads to the systemic stimulation of the DAF-2/IIS pathway and the inhibition of cytoprotective transcription factors, such as DAF-16/FOXO and HSF-1. Here we find that the tyramine-dependent inactivation of DAF-16 and HSF-1 relies on the intestinal release of the ILP INS-3. The tyramine-mediated secretion of INS-3, provides a novel molecular brain-gut communication pathway that links the flight response with the suppression of cytoprotective mechanisms. We show that the release of a neuronal stress neurohormone stimulates the secretion of INS-3 from the intestine to systemically inhibit DAF-2-dependent cytoprotective mechanisms. Importantly, mutants in *ins-3* exhibit greater resistance to environmental stressors compared to the wild-type, even in the absence of tyramine exposure or an active escape response. This enhanced resistance is associated with the detection of a tonic release of this peptide. This finding aligns with the increased resistance to environmental stressors in tyramine-deficient mutants compared to the wild-type. Together, these results imply the existence of a basal release of tyramine that, in turn, activates a tonic level of INS-3 release, partially inhibiting cytoprotective mechanisms. This entire process is further exacerbated during the escape response, where tyramine levels rise.

The secretion polarity of intestinal INS-35 and INS-7 has been demonstrated to be crucial for their function [17]. During reproductive growth, these ILPs are secreted through the basolateral side of the intestinal cell into the pseudocoelomic cavity to inhibit dauer formation [17]. In contrast, in the dauer stage, there is a shift in the polarity of their secretion, and they are released through the apical membrane into the intestinal lumen, where they undergo degradation. The accumulation of INS-3::VENUS in coelomocytes during the flight response suggests that the release of INS-3 occurs on the basolateral side of the intestinal epithelium. It remains to be determined whether tyramine induces a switch in INS-3 secretion polarity, similar to INS-35 and INS-7 during reproductive growth.

Although we show that TYRA-3 stimulates the release of INS-3 in the intestine, the intracellular mechanisms that link these processes are still unknown. ILPs are released from dense core vesicles (DCVs). We have previously shown that the intestinal expression of HID-1, a protein that plays a key role in DCV biogenesis and ILP release in worms and mice [23,51], is required for the tyraminergic inhibition of the stress response [27]. Phylogenetic analyses indicate that TYRA-3 clusters with other GPCRs that are coupled to Gq signaling pathways [52]. Gq signaling stimulates diacylglycerol (DAG) and inositol trisphosphate (IP3) production. Interestingly, studies in mammals have shown that the activation of Gq receptors is essential for insulin secretion, and this process relies on DAG and IP3-dependent mechanisms [53,54]. These findings suggest a potential parallel between the Gq-mediated pathways involved in insulin secretion in mammals and the TYRA-3 signaling pathway in *C. elegans*. Additional experiments are required to unravel the molecular mechanisms by which the activation of a Gq-coupled receptor leads to the release of DCVs containing ILPs.

Although the activation of cytoprotective pathways is crucial to protect against the potential costs of cellular damage, repair, and death, it comes at the expense of resources that could otherwise be allocated to animal development and reproduction [55,56] This is consistent with our observations that stress-resistant *ins-3* mutants, exhibit a slower developmental rate and a reduced brood size. Therefore, it is advantageous for an animal to limit the activation of cytoprotective pathways under favorable conditions, while still retaining the capacity to induce these pathways in emergency situations. This delicate balance between maintaining normal reproductive development while preserving inducible cytoprotective capacity is crucial for the survival and overall fitness of an organism. The neural perception of stressors and subsequent regulation of cytoprotective responses play a pivotal role in this plasticity, ensuring that resources are allocated optimally to support both growth and defense mechanisms.

In vertebrates, the release of adrenaline (the vertebrate counterpart of tyramine) in response to acute stressors is a fundamental component of the flight response [57,58]. Frequent exposure to acute stressors can lead to chronic activation of the stress hormone system [59,60]. Similar to our findings in *C. elegans*, this chronic activation weakens the animal’s defense systems against other stressors, compromising its ability to cope with subsequent challenges [61,62]. In humans, persistent exposure to real or perceived danger, as seen in conditions like Post-Traumatic Stress Disorders (PTSDs), has been associated with adverse health outcomes such as type 2 diabetes, increased oxidative stress, neurodegenerative disorders and premature death [63–68]. It is worth noting that individuals with PTSD often exhibit hyperinsulinemia [69–72] emphasizing the potential role of insulin dysregulation in stress-related health impairments. The precise mechanisms by which increased levels of stress neurohormones impair general health and defense mechanisms in humans are still not fully understood. However, considering the remarkable conservation of neural control over stress responses across species, it would be interesting to investigate whether chronic release of adrenaline negatively impact health by directly stimulating the secretion of ILPs.

## MATERIALS AND METHODS

### Strains and maintenance of C. elegans

Standard *C. elegans* culture and molecular biology methods were used [73,74]. All *C. elegans* strains were grown at 20°C on Nematode Growth Media (NGM) agar plates with OP50 *Escherichia coli* as a food source. Low population density was maintained throughout development and during the assays. The wild type reference strain used in this study is N2 Bristol. Some of the strains were obtained through the Caenorhabditis Genetics Center (CGC, University of Minnesota), which is funded by NIH Office of Research Infrastructure Programs (P40 OD010440). All strains used in this study were backcrossed four times with N2 to eliminate possible background mutations.

The strains used were:

N2 (Wild Type)

OAR186 RB1915 *ins-3 (ok2488)* II 4xBC

MT10661 *tdc-1(n3420)* II

CX11839 *tyra-3(ok325)* X

HT1690 *unc-119(ed3)* III*; wwls26 [Pins-3::GFP + unc-119(+)]*

HT1693 *unc-119(ed3)* III*; wwEx63 [Pins-4::GFP + unc-119(+)]*

OAR45 *nbaEx8 [Pins-3::ins-3 (20)+lin-15 (80)];ins-3 (ok2488), lin-15 (n765ts)]*

OAR43 *nbaEx6 [Pelt-2::ins-3 (20)+lin-15 (80)];ins-3 (ok2488), lin-15 (n765ts)]*

OAR50 *nbaEx13 [Prgef-1::ins-3 (20)+lin-15 (80)];ins-3 (ok2488), lin-15 (n765ts)]*

OAR11 *tdc-1 (n3420); ins-3 (ok2488)*

OAR169 *tyra-3 (n3420); ins-3 (ok2488)*

OAR68 *nbaEx6 [Pelt-2::ins-3 (20)+lin-15 (80)]; tyra-3(ok325)* X*; ins-3 (ok2488)* II*; lin-15 (n765ts)]*

OAR69 *nbaEx6 [Pelt-2::ins-3 (20)+lin-15 (80)]; tdc-1(n3420)* II*; ins-3 (ok2488)* II*; lin-15 (n765ts)]*

QW2436 *Pges-1::ins-3::venus*

TJ356 *zIs356[Pdaf-16::DAF-16a/b::GFP + pRF4]*

OAR58 *zIs356[Pdaf-16::DAF-16a/b::GFP + pRF4]; ins-3 (ok2488)* II

OAR73 *zIs356[Pdaf-16::DAF-16a/b::GFP + pRF4]; nbaEx6[Pelt-2::ins-3 (20)+lin-15 (80)]; ins-3 (ok2488) II; lin-15 (n765ts)]*

CL2070 *dvIs70[Phsp-16.2::GFP + pRF4]*

OAR100 *dvIs70[Phsp-16.2::GFP + pRF4]; ins-3 (ok2488)* II

OAR158 *dvIs70[Phsp-16.2::GFP + pRF4]; ins-3 (ok2488)* II*; nbaEx6[Pelt-2::ins-3 (20)+lin-15 (80)];ins-3 (ok2488), lin-15 (n765ts)*

### RNAi silencing

RNAi was carried out by feeding nematodes with dsRNA-producing bacteria as described previously [75]. RNAi clones that target *C. elegans* insulin genes were obtained from Ahringeŕs feeding RNAi library [76]. RNAi plates were prepared with standard NGM agar supplemented with 25 mg/ml ampicilin and 1 mM isopropyl b-D-1-thiogalactopyranoside (IPTG), poured into 35mm plates and allowed to dry for 7 days at 4°C. Fresh HT115 *E. coli* bacteria transformed with the L4440 empty vector or carrying the appropriate RNAi clone were grown in LB containing 100 mg/ml ampicilin at 37°C overnight. One day before use, 75 µl of concentrated bacteria were seeded and plates were kept overnight for induction of RNAi bacteria.

Animals in the fourth larval stage (L4) were transferred to each plate and maintained at 20°C. Once they laid eggs these animals were removed. Therefore, oxidative stress assays were performed in animals (L4 stage) hatched and grown in RNAi plates.

### Stress resistance assays

#### Oxidative stress

Iron sulfate (FeSO_4_) was used as an oxidative stressor as previously described [27,77]. 20 L4 animals were transferred to 35-mm agar plates (4-5 plates) containing FeSO_4_ at the indicated concentration and time (1 mM, 2 hrs for microscopy and 15 mM, 1h for survival assays). To analyze the effect of tyramine on stress resistance, L4 animals were transferred to four 35-mm NGM agar plates (containing 30-40 animals each) seeded with OP50 bacteria with or without 15 mM exogenous tyramine. 14 hrs later, the stress assays were performed.

#### Heat stress

Thermotolerance assays were performed as described [27]. For each strain, four 35-mm NGM agar plates containing 20 animals (with or without 15 mM exogenous tyramine) were incubated at 35°C for 4 hrs. To ensure proper heat transfer, 6-mm-thick NGM agar plates were used. Animals were synchronized as L4s and used 14 hrs later. Plates were sealed with Parafilm in zip-lock bags, and immersed in a water bath equilibrated to 35°C. Surviving animals were counted after 20 h recovery at 20°C. For all assays, animals were scored as dead if they failed to respond to prodding with a platinum-wire pick to the nose.

### Microscopy and image analysis

For microscopy, age-synchronized animals were mounted in M9 with levamisole (10 mM) onto slides with 2% agarose pads *ins-3* expression pattern was analyzed by confocal microscopy as previously described [27]. Images were acquired on confocal microscopy (LSCM; Leica DMIRE2) with 20X and 63X objectives. For *ins-3 and ins-4* expression levels analysis, animals containing the corresponding transcriptional GFP reporter (*Pins-3::GFP* and *Pins-4:GFP,* respectively) were imaged using an epifluorescence microscope (Nikon Eclipse E-600) coupled to a CCD camera (Nikon K2E Apogee) with 20X objective. Fluorescence intensity was quantified using Image J FIJI software.

### Tyramine supplementation

The assays were performed as described. Tyramine hydrochloride (Alfa Aesar) stocks were made with MilliQ sterile water, and diluted to 15 mM into NGM agar before pouring. Plates were stored at 4°C and used within one week after pouring

### INS-3 release quantification

A population of animals expressing the *Pges-1*::*ins-3::venus* transgene was exposed to tyramine (15 mM, 14 hrs). Individuals were immediately mounted and anesthetized as described above. Images were acquired on confocal microscopy (LSCM; Leica DMIRE2). Images were loaded into ImageJ and the soma of the coelomocytes were identified from their anatomical location. A circular region of interest (ROI) was placed around the coelomocyte to measure the mean fluorescence intensity value of each cell. The values obtained for each condition were normalized to the average intensity of the coelomocytes in the control condition.

### Subcellular DAF-16 localization

DAF-16 translocation was analyzed using strains containing the translational P*daf-16::daf-16a/b::GFP* reporter in a wild type, or insulin mutant background (P*daf-16::daf-16/b::GFP*;*ins-3*; P*daf-16*::*daf-16a/b::GFP*;*ins-3;* P*elt-2*::ins-3). Young adult animals (at least 25 animals per experiment, repeated 3-4 times) in basal conditions or exposed to either mild (35°C, 10 min) or strong heat stress (35°C, 30 min), with or without exogenous tyramine, were mounted to analyze DAF-16 cellular distribution under a fluorescence microscope. The number of GFP-labeled nuclei per animal was quantified using Image J FIJI software and normalized to the naive condition within the day.

### Expression analysis of DAF-2–IIS target genes

*hsp-16.*2 expression levels were analyzed in transgenic strains containing transcriptional reporters, in wild type and insulin mutant backgrounds. Animals were imaged using an epifluorescence microscope (Nikon Eclipse E-600) coupled to a CCD camera (Nikon K2E Apogee). Fluorescence intensity was quantified using Image J FIJI software.

### Molecular biology and cloning

Standard molecular biology techniques were employed for the endogenous rescue of *ins-3.* The entire *ins-3* gene, along with a 2.1 kb upstream region from the start codon, was amplified from genomic DNA. Subsequently, this fragment was cloned into a vector backbone derived from the plasmid Ppd95.75, utilizing the *unc-54* 3’UTR. For the construction of intestinal and neuronal rescue constructs, the *ins-3* DNA (including its intron) from this construct was subcloned behind the intestinal reporters elt-2 and the pan-neuronal *rgef-1* promoter, respectively. Additionally, the *ins-3* gene was subcloned into a plasmid containing the intestinal promoter *ges-1* and the YFP variant Venus [36] to generate the reporter *Pges-1::ins-3::venus*.

The sequences of the constructed plasmids are available on the Open Science Framework platform (https://osf.io/wfgvs/).

Transgenic strains were generated through the microinjection of plasmid DNA at 20 ng/μl into the germline, along with the co-injection marker *lin-15* rescuing plasmid pL15EK (80 ng/μl), into *lin-15(n765ts)* mutant animals. At least three independent transgenic lines were established, and the data presented are from a single representative line.

Multiple mutants were obtained using standard techniques [78]. In brief, 10 males from one of the null mutant strains of interest (*tdc-1* or *tyra-3*) were mated with 2 *ins-3* null mutant hermaphrodites. After 72 hrs, F1 worms (one per plate) were isolated to allow for the production of offspring. The resulting F2 generation was isolated again. The double mutant was identified through PCR analysis of genomic DNA

## Data collection and statistics

All data are represented in a format that shows the distribution (Dot plots) and all the graph elements (median, error bars, etc) are defined in each figure legend. For most of our experiments, as is usual in *C. elegans* research, we used a large number of animals per condition in each assay (typically more than 40–50 animals). This number of animals is large enough to ensure appropriate statistical power in the test used. All the statistical tests were performed after checking normality. Grubbs’test was used for outliers analysis (p<0.05). We performed the experiments at least 3–4 times to ensure reproducibility. All the animals were grown in similar conditions and the experiments were performed on different days, with different animal batches. In general, the experimenter was blind to the conditions/strains tested. Drugs were previously controlled by analyzing a known phenotype (e.g. worm paralysis and head relaxation on tyramine 30mM). Animals used were age-synchronized (L4 or 14h past L4).

## Acknowledgments

Some strains were provided by the CGC, which is funded by NIH Office of Research Infrastructure Programs (P40 OD010440). We thank Jeremy Florman and Andrés Garelli for helpful discussions. In addition, we would like to acknowledge Ignacio Bergé, Andrea Thackeray, Adrian Bizet, Carolina Gomila, Marta Stulhdreher and Carla Chrestía for technical support. We thank Marian Walhout for strains.

---

**Fig S1:**
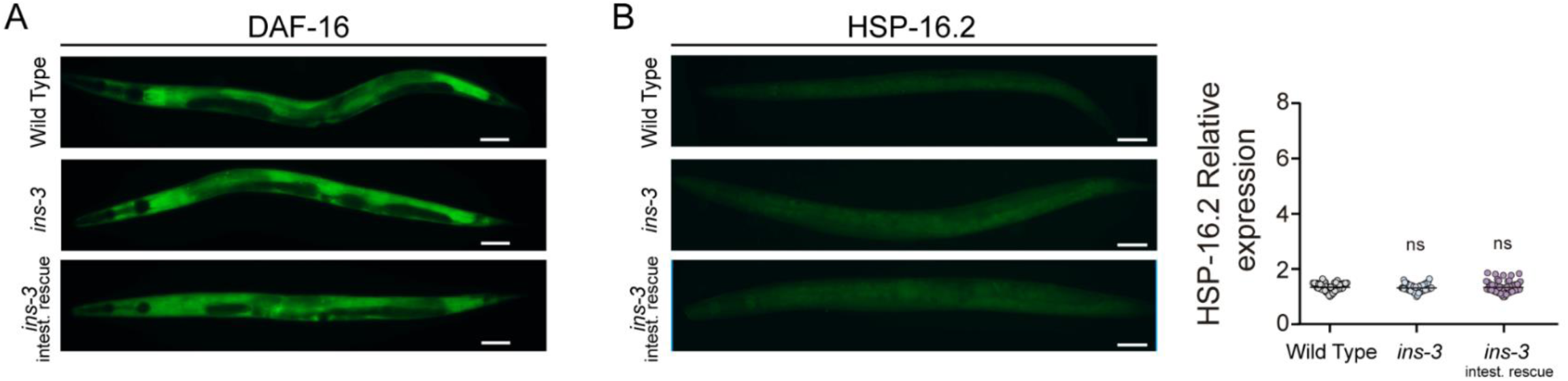
No DAF-16 nuclear translocation or HSP-16.2 induction was observed in non-stressed *ins-3* null mutants. A) Representative fluorescence images (20X) depicting the localization of DAF-16a/b::GFP under basal conditions in WT, *ins-3* null mutant, and intestinal rescue of *ins-3* backgrounds. No nuclear localization of DAF-16 was observed in 20-30 animals for each condition. Scale bar, 50 μm. B) Left. Representative fluorescence images (20X) of animals expressing *Phsp16.2::GFP* in WT, *ins-3* null mutant, and intestinal rescue of *ins-3* backgrounds under basal conditions. Scale bar, 50 μm. Right. Corresponding quantification of the fluorescence level per animal. Scatter dot plot with relative expression of *Phsp16.2::GFP* (normalized to naive animals, line shows median, n= 55-80 per condition). One-way ANOVA and Dunnett’s post-hoc test versus naïve. ns: not significant

**Fig S2:**
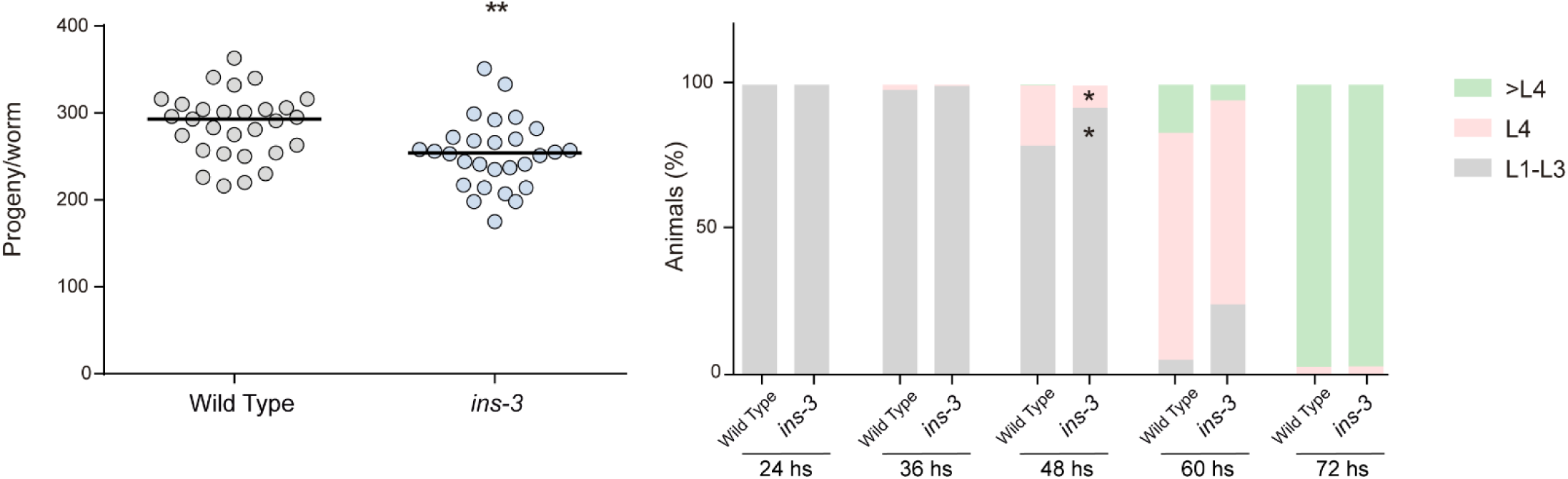
*ins-3* mutants produce reduced offspring and develop more slowly than wild-type animals. Left. Total number of progeny per worm in wild Type and *ins-3* null mutants (n= 25-30 per condition). Two-tailed t-test was used. ** p < 0.01. Right. Developmental rate of wild type and *ins-3* null mutant worms. A color code was used to represent each of the following animal stages: L1–L3: early larval stages, L4: last larval stage, > L4: adult stage. The animal classification was evaluated at the indicated time points (24, 36, 48, 60 and 72 hrs), (n= 5-6, 100-200 animals per condition per experiment). Data are represented in a stacked bar chart as mean. Two-tailed t-test was used. * p < 0.05

**Fig S3:**
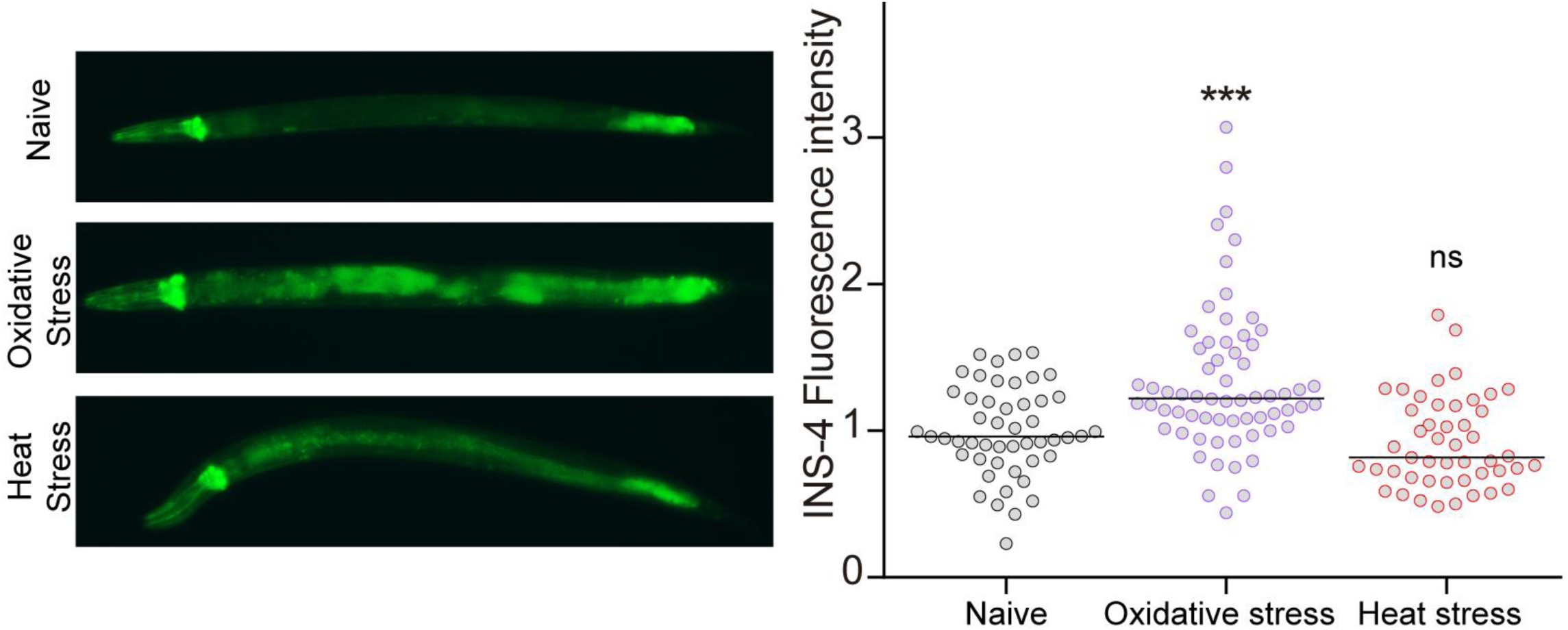
*ins-4* expression is enhanced upon oxidative stress. Left. Representative epifluorescence images (20 X) of young adults expressing Pins-4::GFP under basal conditions, oxidative (1 mM FeSO4 for 2 hrs) or heat stress (30°C for 6 hrs). Scale bar, 50 µm. Right. Corresponding quantification of fluorescence levels per worm. Scatter dot plot with relative expression of *Pins-4::GFP* normalized to naive animals. Line at the median. n= 45-60 per condition. One-way ANOVA (Kruskal–Wallis test) and Dunn’s post-hoc test versus naïve was used. ns: not significant, *** p<0.001

**Fig S4.**
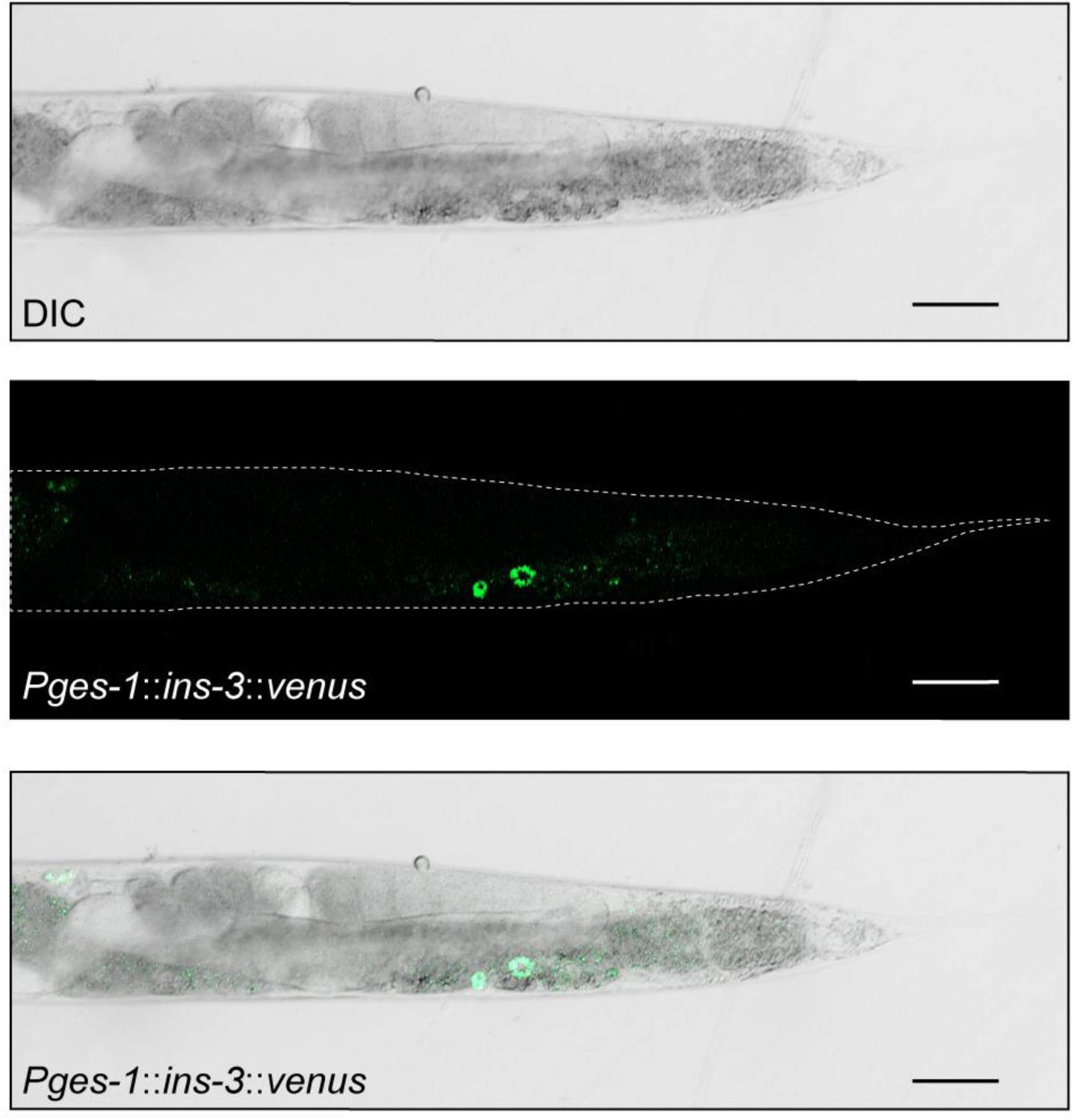
Under basal conditions, INS-3 is tonically released from the intestine. Representative image (40X) of L4-staged worm carrying P*ges-1*::*ins-3*::*venus* in basal conditions showed on DIC, fluorescence, and merged. Note the fluorescence both in the intestine (faint) and in the coelomocytes. Scale bar, 50 μm

**Fig S5.**
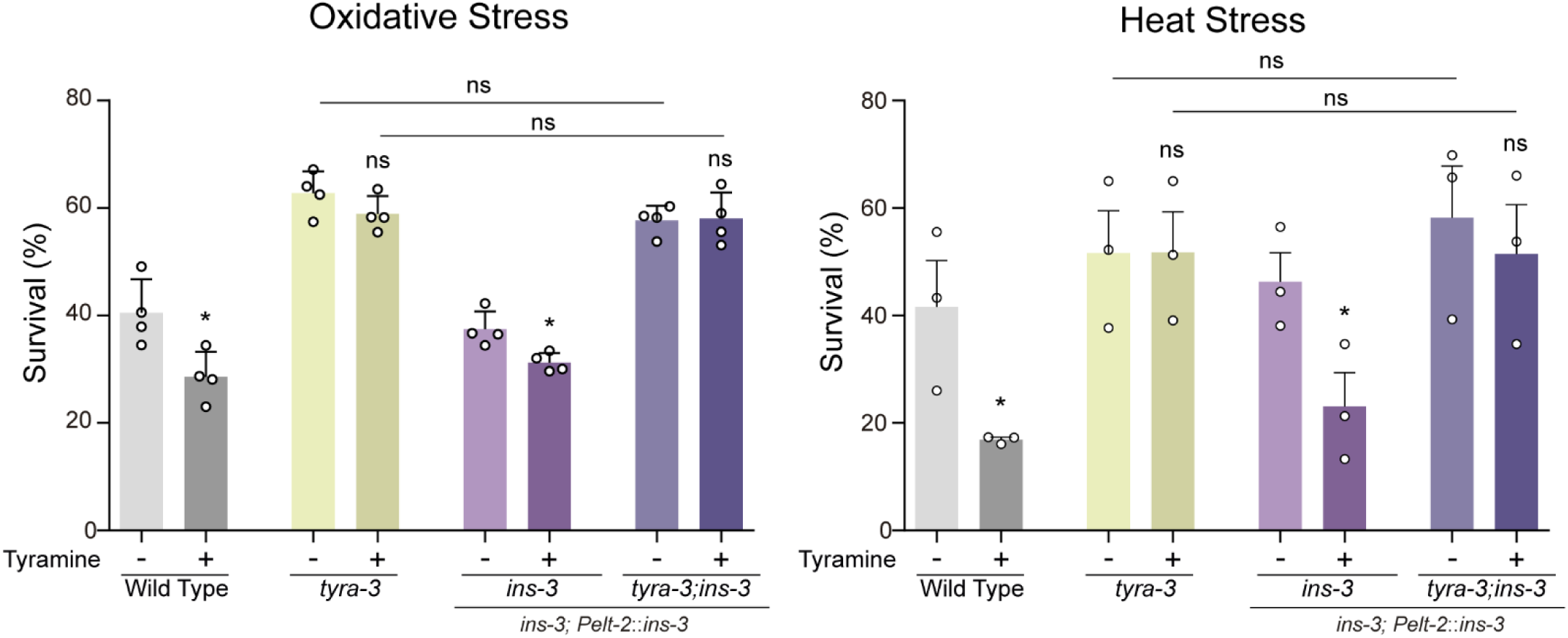
Tyraminergic modulation of stress resistance through intestinal INS-3 release requires the GPCR TYRA-3. Survival percentages to oxidative (left) and heat stress (right) of wild-type, *tyra-3* null mutants, and animals expressing the intestinal rescue of *ins-3* on wild-type and *tyra-3* null mutant backgrounds in the absence or presence of exogenous tyramine (15 mM), (mean ± s.e.m., n= 3-4, 40-50 animals per condition). For conditions with tyramine, two-tailed Student’s t-test versus same strain without tyramine was used. ns: not significant, * p<0.05.

## Reference List

1. Rossnerova A, Izzotti A, Pulliero A, Bast A (2020) The Molecular Mechanisms of Adaptive Response Related to Environmental Stress. 21.

2. Ellis BJ, Jackson JJ, Boyce WT (2006) The stress response systems: Universality and adaptive individual differences. Developmental Review 26: 175–212.

3. Nussey DH, Wilson AJ, Brommer JE (2007) The evolutionary ecology of individual phenotypic plasticity in wild populations. J Evol Biol 20: 831–844.

4. Auer SK, Salin K, Rudolf AM, Anderson GJ, Metcalfe NB (2015) Flexibility in metabolic rate confers a growth advantage under changing food availability. J Anim Ecol 84: 1405–1411.

5. Zhang N, Cao L (2017) Starvation signals in yeast are integrated to coordinate metabolic reprogramming and stress response to ensure longevity. Curr Genet 63: 839–843.

6. Moreau K, Luo S, Rubinsztein DC (2010) Cytoprotective roles for autophagy. Curr Opin Cell Biol 22: 206–211.

7. Murphy CT, Hu PJ (2013) Insulin/insulin-like growth factor signaling in C. elegans. WormBook: 1–43.

8. Biglou SG, Bendena WG, Chin-Sang I (2021) An overview of the insulin signaling pathway in model organisms Drosophila melanogaster and Caenorhabditis elegans. Peptides 145: 170640.

9. Kenyon CJ (2010) The genetics of ageing. Nature 464: 504–512.

10. Lithgow GJ, White TM, Melov S, Johnson TE (1995) Thermotolerance and extended life-span conferred by single-gene mutations and induced by thermal stress. Proc Natl Acad Sci U S A 92: 7540–7544.

11. Honda Y, Honda S (1999) The daf-2 gene network for longevity regulates oxidative stress resistance and Mn-superoxide dismutase gene expression in Caenorhabditis elegans. FASEB J 13: 1385–1393.

12. Kenyon C, Chang J, Gensch E, Rudner A, Tabtiang R (1993) A C. elegans mutant that lives twice as long as wild type. Nature 366: 461–464.

13. Lithgow GJ, White TM, Hinerfeld DA, Johnson TE (1994) Thermotolerance of a long-lived mutant of Caenorhabditis elegans. J Gerontol 49: B270–B276.

14. Pierce SB, Costa M, Wisotzkey R, Devadhar S, Homburger SA, et al. (2001) Regulation of DAF-2 receptor signaling by human insulin and ins-1, a member of the unusually large and diverse C. elegans insulin gene family. Genes & development Genes Dev. pp. 672–686.

15. Chen Z, Hendricks M, Cornils A, Maier W, Alcedo J, et al. (2013) Two insulin-like peptides antagonistically regulate aversive olfactory learning in C. elegans. Neuron 77: 572–585.

16. Matsunaga Y, Nakajima K, Gengyo-Ando K, Mitani S, Iwasaki T, et al. (2012) A Caenorhabditis elegans insulin-like peptide, INS-17: its physiological function and expression pattern. Biosci Biotechnol Biochem 76: 2168-2172.

17. Matsunaga Y, Honda Y, Honda S, Iwasaki T, Qadota H, et al. (2016) Diapause is associated with a change in the polarity of secretion of insulin-like peptides. Nat Commun 7: 10573.

18. Michaelson D, Korta DZ, Capua Y, Hubbard EJ (2010) Insulin signaling promotes germline proliferation in C. elegans. Development 137: 671–680.

19. Ritter AD, Shen Y, Fuxman BJ, Jeyaraj S, Deplancke B, et al. (2013) Complex expression dynamics and robustness in C. elegans insulin networks. Genome Res 23: 954–965.

20. Zheng S, Chiu H, Boudreau J, Papanicolaou T, Bendena W, et al. (2018) A functional study of all 40 Caenorhabditis elegans insulin-like peptides. J Biol Chem 293: 16912–16922.

21. Kao G, Nordenson C, Still M, Ronnlund A, Tuck S, et al. (2007) ASNA-1 positively regulates insulin secretion in C. elegans and mammalian cells. Cell 128: 577–587.

22. Yoshina S, Mitani S (2015) Loss of C. elegans GON-1, an ADAMTS9 Homolog, Decreases Secretion Resulting in Altered Lifespan and Dauer Formation. PLoS One 10: e0133966.

23. Mesa R, Luo S, Hoover CM, Miller K, Minniti A, et al. (2011) HID-1, a new component of the peptidergic signaling pathway. Genetics 187: 467–483.

24. Arun CP (2004) Fight or flight, forbearance and fortitude: the spectrum of actions of the catecholamines and their cousins. Ann N Y Acad Sci 1018: 137–140.

25. McCarty R (2016) Chapter 4 - The Fight-or-Flight Response: A Cornerstone of Stress Research. In: Fink G, editor. Stress: Concepts, Cognition, Emotion, and Behavior. San Diego: Academic Press. pp. 33–37.

26. Mronz M, Lehmann FO (2008) The free-flight response of Drosophila to motion of the visual environment. J Exp Biol 211: 2026–2045.

27. De Rosa MJ, Veuthey T, Florman J, Grant J, Blanco MG, et al. (2019) The flight response impairs cytoprotective mechanisms by activating the insulin pathway. Nature 573: 135–138.

28. Alkema MJ, Hunter-Ensor M, Ringstad N, Horvitz HR (2005) Tyramine Functions independently of octopamine in the Caenorhabditis elegans nervous system. Neuron 46: 247–260.

29. Donnelly JL, Clark CM, Leifer AM, Pirri JK, Haburcak M, et al. (2013) Monoaminergic orchestration of motor programs in a complex C. elegans behavior. PLoS Biol 11: e1001529.

30. Maguire SM, Clark CM, Nunnari J, Pirri JK, Alkema MJ (2011) The C. elegans touch response facilitates escape from predacious fungi. Curr Biol 21: 1326–1330.

31. Pirri JK, McPherson AD, Donnelly JL, Francis MM, Alkema MJ (2009) A tyramine-gated chloride channel coordinates distinct motor programs of a Caenorhabditis elegans escape response. Neuron 62: 526–538.

32. Pirri JK, Rayes D, Alkema MJ (2015) A Change in the Ion Selectivity of Ligand-Gated Ion Channels Provides a Mechanism to Switch Behavior. PLoS Biol 13: e1002238.

33. Chiang WC, Ching TT, Lee HC, Mousigian C, Hsu AL (2012) HSF-1 regulators DDL-1/2 link insulin-like signaling to heat-shock responses and modulation of longevity. Cell 148: 322–334.

34. Zhang YP, Zhang WH, Zhang P, Li Q, Sun Y, et al. (2022) Intestine-specific removal of DAF-2 nearly doubles lifespan in Caenorhabditis elegans with little fitness cost. Nat Commun 13: 6339.

35. Lee SH, Omi S, Thakur N, Taffoni C, Belougne J, et al. (2018) Modulatory upregulation of an insulin peptide gene by different pathogens in C. elegans. 9: 648–658.

36. Florman JT, Alkema MJ (2022) Co-transmission of neuropeptides and monoamines choreograph the C. elegans escape response. 18: e1010091.

37. Mahoney TR, Luo S, Round EK, Brauner M, Gottschalk A, et al. (2008) Intestinal signaling to GABAergic neurons regulates a rhythmic behavior in Caenorhabditis elegans. Proc Natl Acad Sci U S A 105: 16350–16355.

38. Baumeister R, Schaffitzel E, Hertweck M (2006) Endocrine signaling in Caenorhabditis elegans controls stress response and longevity. J Endocrinol 190: 191–202.

39. Scott BA, Avidan MS, Crowder CM (2002) Regulation of hypoxic death in C. elegans by the insulin/IGF receptor homolog DAF-2. Science 296: 2388–2391.

40. Barsyte D, Lovejoy DA, Lithgow GJ (2001) Longevity and heavy metal resistance in daf-2 and age-1 long-lived mutants of Caenorhabditis elegans. FASEB J 15: 627–634.

41. Vanfleteren JR (1993) Oxidative stress and ageing in Caenorhabditis elegans. Biochem J 292 (Pt 2): 605–608.

42. Li C, Kim K (2008) Neuropeptides. WormBook: 1–36.

43. Ewe CK, Alok G, Rothman JH (2021) Stressful development: integrating endoderm development, stress, and longevity. Developmental Biology 471: 34–48.

44. Uno M, Tani Y, Nono M, Okabe E, Kishimoto S, et al. (2021) Neuronal DAF-16-to-intestinal DAF-16 communication underlies organismal lifespan extension in C. elegans. iScience 24: 102706.

45. Apfeld J, Kenyon C (1999) Regulation of lifespan by sensory perception in Caenorhabditis elegans. Nature 402: 804–809.

46. Wolkow CA, Kimura KD, Lee MS, Ruvkun G (2000) Regulation of C. elegans life-span by insulinlike signaling in the nervous system. Science 290: 147–150.

47. Iser WB, Gami MS, Wolkow CA (2007) Insulin signaling in Caenorhabditis elegans regulates both endocrine-like and cell-autonomous outputs. Dev Biol 303: 434–447.

48. Suzuki A, Nakauchi H, Taniguchi H (2003) Glucagon-like peptide 1 (1-37) converts intestinal epithelial cells into insulin-producing cells. Proc Natl Acad Sci U S A 100: 5034–5039.

49. Talchai C, Xuan S, Kitamura T, DePinho RA, Accili D (2012) Generation of functional insulin-producing cells in the gut by Foxo1 ablation. Nat Genet 44: 406–412, S401.

50. Srivastava S, Pandey H, Singh SK, Tripathi YB (2019) GLP 1 Regulated Intestinal Cell’s Insulin Expression and Selfadaptation before the Onset of Type 2 Diabetes. Adv Pharm Bull 9: 325–330.

51. Du W, Zhou M, Zhao W, Cheng D, Wang L, et al. (2016) HID-1 is required for homotypic fusion of immature secretory granules during maturation. Elife 5.

52. Wragg RT, Hapiak V, Miller SB, Harris GP, Gray J, et al. (2007) Tyramine and octopamine independently inhibit serotonin-stimulated aversive behaviors in Caenorhabditis elegans through two novel amine receptors. J Neurosci 27: 13402–13412.

53. Latour MG, Alquier T, Oseid E, Tremblay C, Jetton TL, et al. (2007) GPR40 is necessary but not sufficient for fatty acid stimulation of insulin secretion in vivo. Diabetes 56: 1087–1094.

54. Sassmann A, Gier B, Grone HJ, Drews G, Offermanns S, et al. (2010) The Gq/G11-mediated signaling pathway is critical for autocrine potentiation of insulin secretion in mice. J Clin Invest 120: 2184–2193.

55. Crawford D, Libina N, Kenyon C (2007) Caenorhabditis elegans integrates food and reproductive signals in lifespan determination. Aging Cell 6: 715–721.

56. Swanson MM, Riddle DL (1981) Critical periods in the development of the Caenorhabditis elegans dauer larva. Dev Biol 84: 27–40.

57. Cannon WB (1929) ORGANIZATION FOR PHYSIOLOGICAL HOMEOSTASIS. Physiological Reviews 9: 399–431.

58. Selye H (1974) Stress without distress: Philadelphia; New York: J.B. Lippincott.

59. Boonstra R (2013) Reality as the leading cause of stress: rethinking the impact of chronic stress in nature. Functional Ecology 27: 13.

60. Sabban EL, Kvetnansky R (2001) Stress-triggered activation of gene expression in catecholaminergic systems: dynamics of transcriptional events. Trends Neurosci 24: 91–98.

61. Odio M, Brodish A, Ricardo MJ, Jr. (1987) Effects on immune responses by chronic stress are modulated by aging. Brain Behav Immun 1: 204–215.

62. Travers M, Clinchy M, Zanette L, Boonstra R, Williams TD (2010) Indirect predator effects on clutch size and the cost of egg production. Ecol Lett 13: 980–988.

63. Aamodt EJ, Chung MA, McGhee JD (1991) Spatial control of gut-specific gene expression during Caenorhabditis elegans development. Science 252: 579–582.

64. Dhabhar FS (2014) Effects of stress on immune function: the good, the bad, and the beautiful. Immunol Res 58: 193–210.

65. Glaser R, Kiecolt-Glaser J (2005) How stress damages immune system and health. Discov Med 5: 165–169.

66. Groer MW, Kane B, Williams SN, Duffy A (2015) Relationship of PTSD Symptoms With Combat Exposure, Stress, and Inflammation in American Soldiers. Biol Res Nurs 17: 303–310.

67. Miller MW, Sadeh N (2014) Traumatic stress, oxidative stress and post-traumatic stress disorder: neurodegeneration and the accelerated-aging hypothesis. Mol Psychiatry 19: 1156–1162.

68. Porhomayon J, Kolesnikov S, Nader ND (2014) The Impact of Stress Hormones on Post-traumatic Stress Disorders Symptoms and Memory in Cardiac Surgery Patients. J Cardiovasc Thorac Res 6: 79–84.

69. Modan M, Halkin H (1991) Hyperinsulinemia or Increased Sympathetic Drive as Links for Obesity and Hypertension. Diabetes Care 14: 470–487.

70. Mellon SH, Bersani FS, Lindqvist D, Hammamieh R, Donohue D, et al. (2019) Metabolomic analysis of male combat veterans with post traumatic stress disorder. 14: e0213839.

71. Aaseth J, Roer GE, Lien L, Bjørklund G (2019) Is there a relationship between PTSD and complicated obesity? A review of the literature. Biomedicine & Pharmacotherapy 117: 108834.

72. Michopoulos V, Vester A, Neigh G (2016) Posttraumatic stress disorder: A metabolic disorder in disguise? Exp Neurol 284: 220–229.

73. Brenner S (1974) The genetics of Caenorhabditis elegans. Genetics 77: 71–94.

74. Stiernagle T (2006) Maintenance of C. elegans. WormBook: 1–11.

75. Conte D, Jr., MacNeil LT, Walhout AJ, Mello CC (2015) RNA Interference in Caenorhabditis elegans. Curr Protoc Mol Biol 109: 26–30.

76. Kamath RS, Fraser AG, Dong Y, Poulin G, Durbin R, et al. (2003) Systematic functional analysis of the Caenorhabditis elegans genome using RNAi. Nature 421: 231–237.

77. Andersen N, Veuthey T, Blanco MG, Silbestri GF, Rayes D, et al. (2022) 1-Mesityl-3-(3-Sulfonatopropyl) Imidazolium Protects Against Oxidative Stress and Delays Proteotoxicity in C. elegans. Frontiers in Pharmacology 13.

78. Fay DS (2013) Classical genetic methods. WormBook: 1–58.

